# A pH-eQTL interaction at the RIT2-SYT4 Parkinson’s disease risk locus in the substantia nigra

**DOI:** 10.1101/2020.12.16.423140

**Authors:** Sejal Patel, Derek Howard, Leon French

## Abstract

Parkinson’s disease (PD) causes severe motor and cognitive disabilities that result from the progressive loss of dopamine neurons in the substantia nigra. The rs12456492 variant in the RIT2 gene has been repeatedly associated with increased risk for Parkinson’s disease. From a transcriptomic perspective, a meta-analysis found that *RIT2* gene expression is correlated with pH in the human brain. To assess these pH associations in relation to PD risk, we examined the two datasets that assayed rs12456492, gene expression, and pH in the postmortem human brain. Using the BrainEAC dataset, we replicate the positive correlation between *RIT2* gene expression and pH in the human brain (n=100). Furthermore, we found that the relationship between expression and pH is influenced by rs12456492. When tested across ten brain regions, this interaction is specifically found in the substantia nigra. A similar association was found for the co-localized *SYT4* gene. In addition, *SYT4* associations are stronger in a combined model with both genes, and the *SYT4* interaction appears to be specific to males. In the GTEx dataset, the pH associations involving rs12456492 and expression of either *SYT4* and *RIT2* was not seen. This null finding may be due to the short postmortem intervals (PMI) of the GTEx tissue samples. In the BrainEAC data, we tested the effect of PMI and only observed the interactions in the longer PMI samples. These previously unknown associations suggest novel mechanistic roles for rs12456492, *RIT2*, and *SYT4* in the regulation of pH in the substantia nigra.

## Introduction

Parkinson’s disease (PD) is a common neurodegenerative disease characterized by the loss of dopamine neurons in the substantia nigra. Individuals with PD show severe motor and cognitive disabilities. The etiology of PD is complex, with multiple genetic and environmental risk factors (Nalls et al. 2019; Bellou et al. 2016).

Recent GWAS studies of sporadic PD found that common genetic variants explain about 16-36% of PD’s heritable risk (Nalls et al. 2019). Most studies have focused on genes that have been associated with both monogenic and sporadic forms of PD. These include *SNCA, LRRK2*, and *GBA* (Chang et al. 2017). However, over 90 independent risk signals have been identified. These recent GWAS hits are underexplored in the context of PD. For example, rs12456492 was first associated with PD in a 2011 GWAS study (Do et al. 2011). Subsequent studies have replicated this locus on chromosome 18, confirming an association with PD (Nalls et al. 2019). This locus contains the Ras Like Without CAAX 2 (*RIT2*) and Synaptotagmin 4 (*SYT4*) genes (Pankratz et al. 2012). The *SYT4* gene is a member of the synaptotagmin family and regulates synaptic transmission (Dean et al. 2009). In the context of PD, Mendez et al. demonstrated that somatodendritic dopamine release depends on *SYT4* (Mendez et al. 2011). However, the *RIT2* and *SYT4* genes have not been extensively characterized in relation to PD.

In a cross laboratory comparison of expression profiling data from normal human postmortem brains, Mistry and Pavlidis identified a robust correlation between RIT2 and tissue pH (Mistry and Pavlidis 2010). This meta-analysis included 11 studies that provided 421 cortical transcriptomes. In this meta-analysis, RIT2 was ranked tenth on the pH up-regulation list of 15,845 genes. The regulation of pH within the brain is crucial for proper physiological functioning. Specifically, impairment in this regulation can alter neuronal state leading to physiopathological conditions (Sinning and Hübner 2013). Results from the Mistry et al. meta-analysis suggest that RIT2 may be involved in neural pH regulation. Dysregulated pH can lead to oxidative stress and influence alpha-synuclein aggregation, which may play a role in PD pathology (Hwang 2013). These findings motivate a deeper characterization of pH and *RIT2* gene expression in the context of PD.

In this study, we use the BrainEac data to replicate the correlation between *RIT2* expression and pH. In addition, we further test for associations between *SYT4* and pH. We characterize interactions involving a co-localized genetic risk variant for PD, pH, *RIT2* and SYT4 gene expression. We explore these interactions in two independent postmortem brain datasets and test the impact of PMI and sex. We also perform co-expression searches to associate genes of known function to *RIT2* and *SYT4*.

## Methods

### BrainEac dataset

Phenotype, genome-wide expression, and genotype information were obtained from The Brain expression quantitative trait loci (eQTL) Almanac (BrainEAC) project. This data from the UK Brain expression consortium was generated to investigate genetic regulation and alternative splicing. The consortium assayed genome-wide expression in ten brain regions using Affymetrix Exon 1.0 ST arrays (Illumina, San Diego, CA, USA) from 134 neuropathologically normal donors (Ramasamy et al. 2014). We extracted genotype data and expression values for the *RIT2, SYT4* and *CA10* genes from the BrainEAC web-based resource (http://www.braineac.org/). Age, sex, postmortem interval (PMI), RNA integrity number (RIN) and pH data were obtained from Gene Expression Omnibus (GSE46706) (Edgar, Domrachev, and Lash 2002). Of the 134 brains, we restricted our analyses to the 100 brains from the Medical Research Council (MRC) Sudden Death Brain and Tissue Bank in Edinburgh, UK (Millar et al. 2007) that had pH and genotype data. For each brain, pH was measured in the lateral ventricle because it is known to be stable across brain regions (Trabzuni et al. 2012). All ten regions were not sampled in all the brains, resulting in 73 substantia nigra samples. The average age of the cohort was 51.86, with 76.71% composed of males.

### GTEx dataset

The Genotype-Tissue Expression (GTEx) project was designed to identify eQTL’s in diverse human tissues (Lonsdale et al. 2013). We extracted version 8 of GTEx data, which included expression data for the substantia nigra and genotype data for PD risk SNP rs12456492, resulting in a sample size of 113. Tissue samples were collected from non-diseased postmortem brain samples. RNA sequencing expression data was obtained from the GTEx Portal and log normalized (filename: GTEx_Analysis_2017-06-05_v8_RNASeQCv1.1.9_gene_tpm.gct). Genotype information was obtained from whole-exome sequencing by the GTEx consortium (filename:

GTEx_Analysis_2017-06-05_v8_WholeExomeSeq_979Indiv_VEP_annot.vcf). Additional information extracted from the GTEx Portal included age, sex, PMI, RIN and pH (measured in the cerebellum) (filename: GTEx_Analysis_v8_Annotations_SampleAttributesDS.txt). The average age of the cohort was 57.87 years, with 70.8% composed of males.

### Co-expression meta-analysis tools

SEEK (Search-Based Exploration of Expression Compendium for Humans) was used to identify top genes co-expressed with *RIT2*. The top-ranked dataset from the co-expression result was downloaded to test to detail correlations between RIT2, SYT4 and CA10. Normalized expression data for GSE20146 was obtained from the Gemma web-based tool, using the filter option, which resulted in the removal of one outlier (GSM505262) (Lim et al. 2021).

### Statistical analysis

Pearson correlation and ordinary least square linear models were performed in R (Core 2013).

### Availability of Code and Data

Scripts and data for reproducing the majority of the analyses are publicly available online at https://github.com/Sejal24/PD_Manuscript_RIT2_SYT4_pH. The GTEx data is available via DBGap (Accession: phs000424.v8.p2).

## Results

### Brain-wide *RIT2* and *SYT4* gene expression is correlated with pH

To replicate the association between pH and *RIT2* and additionally test *SYT4*, we examined their relationships. The gene expression of *RIT2* and *SYT4* was averaged across all ten brain regions assayed in the BrainEAC study (n=100 brains). We observed a broad range of pH values (5.42 to 6.63). Gene expression and brain pH was correlated for *RIT2* (r = 0.59, p < 0.0001), and *SYT4* (r = 0.58, p < 0.0001). As seen in Figure 1, five brains had abnormally low pH values, with pH values were lower than 2 standard deviations from the mean (ph < 5.8). To prevent these outliers from skewing our downstream results, we have removed them from all subsequent analyses. After removal of these pH outlier brains, *RIT2* remains correlated with pH (r = 0.22, p < 0.04), however *SYT4* was no longer correlated with brain pH (*SYT4*: r = 0.14, p = 0.17). In agreement with Mistry et al., we also observe a correlation between postmortem brain pH and *RIT2* gene expression.

**Figure 1.**
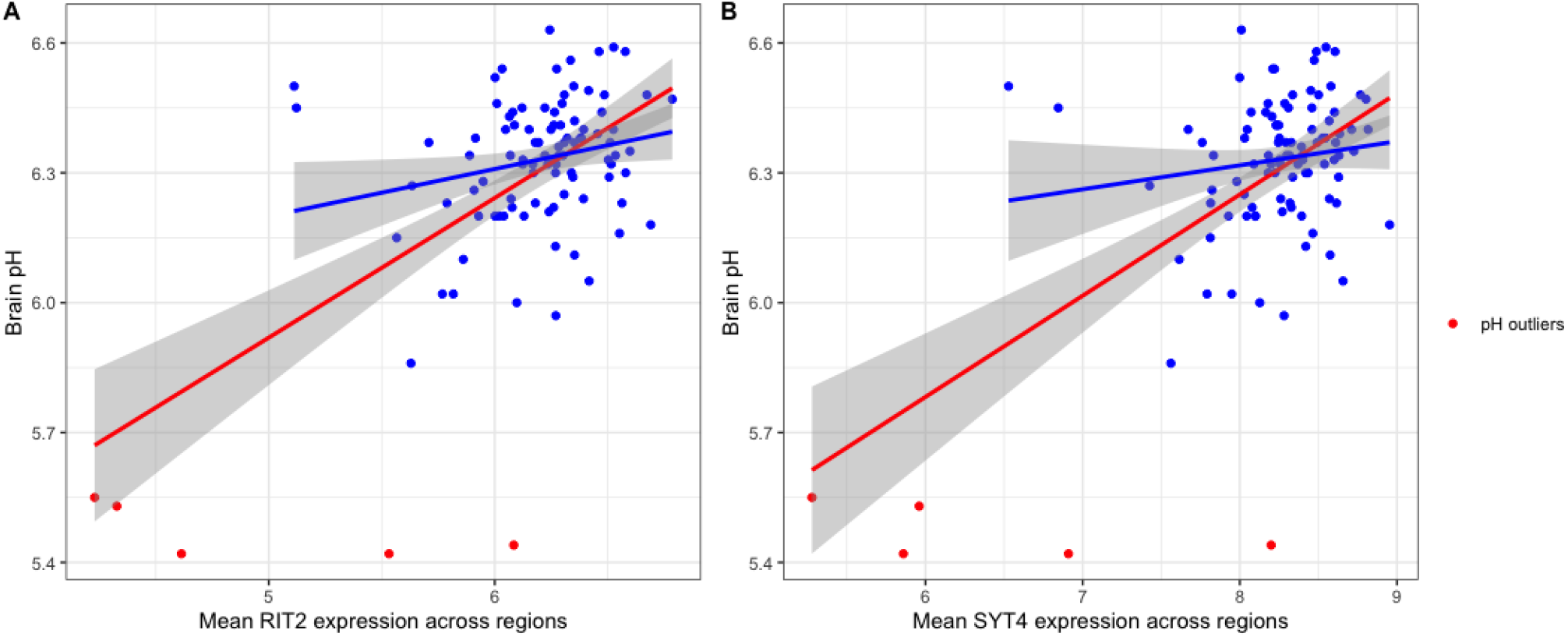
Scatter plots of the relationship between pH with *RIT2* and *SYT4* gene expression. pH outliers are colored red. Lines representing linear fits with (blue) and without outliers (red) include shaded areas marking the 0.95 confidence intervals.

Although pH was measured at the lateral ventricle, it is known that pH is relatively consistent brain-wide (Trabzuni et al. 2011). However, we tested if the correlation with gene expression varies across the ten brain regions profiled. For each individual brain region, *RIT2* was most correlated with pH in the thalamus (n = 90, r = 0.29, p = 0.005, p_FDR_ = 0.055) and substantia nigra (n = 70, r = 0.27, p = 0.026, p_FDR_ = 0.13). In contrast, the correlation between *SYT4* expression and pH was not statistically significant in any of the 10 regions. *SYT4* correlation was high but not significant in the thalamus (r = 0.20, p = 0.057). White matter and regions enriched for white matter (putamen and medulla) had the lowest pH correlations for *RIT2* gene expression, whereas the medulla and substantia nigra had the lowest pH correlation for *SYT4*.

### Rs12456492 influences the association between *RIT2* and *SYT4* expression and pH in the substantia nigra

Given that the single nucleotide polymorphism (SNP) rs12456492 lies between *RIT2* and *SYT4* is associated with Parkinson’s disease, we tested if it is associated with brain pH. Genotype at rs12456492 by itself was associated with brain pH (B = 0.06, p = 0.043). We next tested if this neighbouring PD risk variant influenced the correlations between gene expression and pH. As depicted in Figures 2 and 3, an interaction between pH, rs12456492 and either *RIT2* or *SYT4*, was observed in the substantia nigra (*RIT2*: B = −0.15, p < 0.007, p_FDR_ <0.07, *SYT4*: B = −0.16, p < 0.0001, p_FDR_ < 0.001) but none of the other profiled regions. As shown in Table 1, after accounting for the covariates of sex, age, PMI and RIN, these signals remain. Compared to all other terms in the model, the pH-eQTL interaction for either *RIT2* or *SYT4* expression is the most significant (p<0.005) (Table 1). Specifically, in the BrainEAC data, individuals carrying the risk allele (AG and GG), had a positive correlation between gene expression and pH but not in the AA genotype group. This suggests a weaker coupling between pH, *RIT2* and *SYT4* expression in the substantia nigra may be protective.

**Figure 2.**
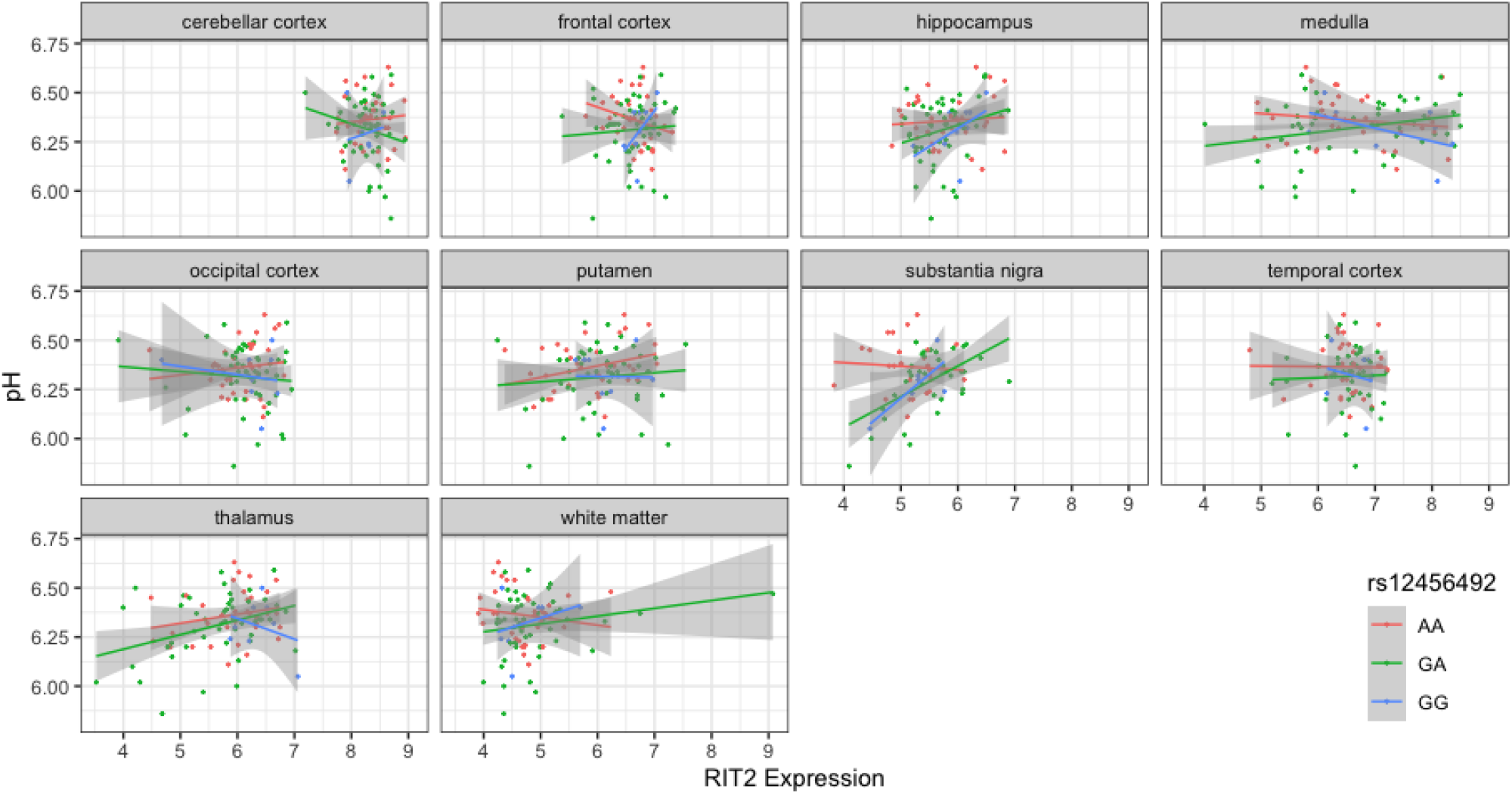
Scatter plot of pH and *RIT2* gene expression based on rs12456492 genotype for 10 brain regions. Genotype groups are colored with lines representing linear fits with shaded areas marking the 0.95 confidence interval.

**Figure 3.**
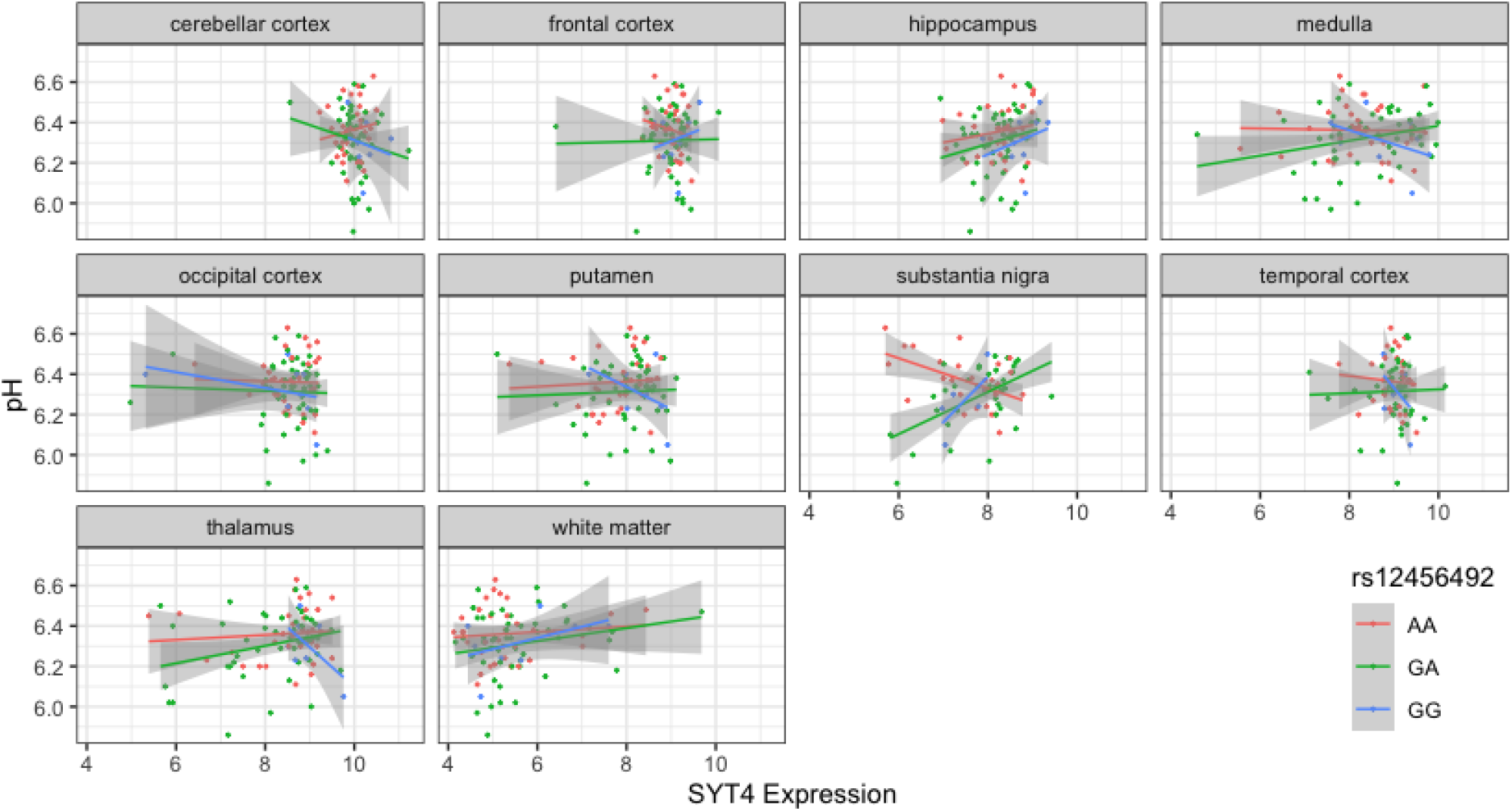
Scatter plot of pH and *SYT4* gene expression based on rs12456492 genotype for 10 brain regions. Genotype groups are colored with lines representing linear fits with shaded areas marking the 0.95 confidence interval.

**Table 1:** Effects of Rs12456492, *RIT2*, and *SYT4* gene expression on brain pH across six models.

|  | Model 1 | Model 2 | Model 3 | Model 4 | Model 5 | Model 6 |
| --- | --- | --- | --- | --- | --- | --- |
| **Rs12456492 (SNP)** | 0.06<br>(0.006-0.12),<br>$p<0.04$ | 0.94<br>(0.35-1.53),<br>$p<0.003$ | 1.38<br>(0.84-1.92),<br>$p<0.00001$ | 0.90<br>(0.31 - 1.49),<br>$p<0.004$ | 1.23<br>(0.71 - 1.77),<br>$p<0.00002$ | 1.23<br>(0.64 - 1.81),<br>$p<0.0001$ |
| **RIT2** | | 0.30<br>(0.14-0.45),<br>$p<0.0004$ | | 0.34<br>(0.18 - 0.51),<br>$p<0.0001$ | 0.11<br>(0.02 - 0.20),<br>$p<0.03$ | 0.10<br>( -0.12 - 0.32),<br>$p<0.37$ |
| **SYT4** | | | 0.26<br>(0.16-0.37),<br>$p<0.00001$ | -0.05<br>( -0.12 - 0.01),<br>$p=0.11$ | 0.17<br>(0.05 - 0.30),<br>$p<0.006$ | 0.18<br>(0.02 - 0.34),<br>$p<0.03$ |
| **RIT2×SNP** | | -0.17<br>(-0.23-<br>-0.05), p<0.005 | | -0.16<br>(-0.27 - -0.04),<br>$p<0.007$ | | 0.01<br>(-0.14 - 0.15),<br>$p=0.93$ |
| **SYT4×SNP** |  |  | -0.17<br>(-0.24 - -0.10),<br>p<0.00001 |  | -0.15<br>(-0.22 - -0.08),<br>p<0.00006 | -0.16<br>(-0.26 - -0.06),<br>p<0.003 |
| **Adjusted R2** | 0.013 | 0.20 | 0.28 | 0.22 | 0.33 | 0.32 |

\* After adjusting for sex, age, PMI and RIN

To investigate the combined influences of *RIT2* and *SYT4* on pH level, we tested three additional models (Table 1, models 4-6). After including all covariates, the addition of *SYT4* gene expression resulted in a slightly better fit in the *RIT2* interaction model (Model 2 vs 4, R^2^ from 0.2 to 0.22, p=0.11). Similarly, the addition of *RIT2* expression explained slightly more variance in the *SYT4* interaction model (Model 3 vs 5, R^2^ from 0.28 to 0.33, p=0.02). Models that have the *SYT4* genetic interaction explain more variance than those that include the *RIT2* interaction. Furthermore, in a model with all tested therms, the *RIT2* genetic interaction term is no longer significant (Model 6).

### Sex-specific signals

To investigate if sex influenced the pH-eQTL interaction we tested a three-way interaction between gene expression, genotype, and gender. This added interaction term was not significant for *RIT2* (B=0.21 [-0.05 - 0.48], p=0.11), but was for *SYT4* (B=0.18 [0.006 - 0.35], p<0.042) when added to Models 2 and 3. We additionally stratified our analyses to test for sex-specific effects on Model 2 and Model 3 (Figure 4). This yielded 54 male and a small group of 16 female samples that lack individuals carrying the GG genotype. In both models, pH-eQTL interactions were significant for males (*RIT2*: B = −0.21, p<0.0001; *SYT4*: B=-0.21, p<0.003) but not females (*RIT2*: B=-0.03, p=0.88; *SYT4*: B = −0.09, p=0.47). Overall, the *SYT4* pH-eQTL appears to be sex-specific, but this analysis is limited by sample size.

**Figure 4:**
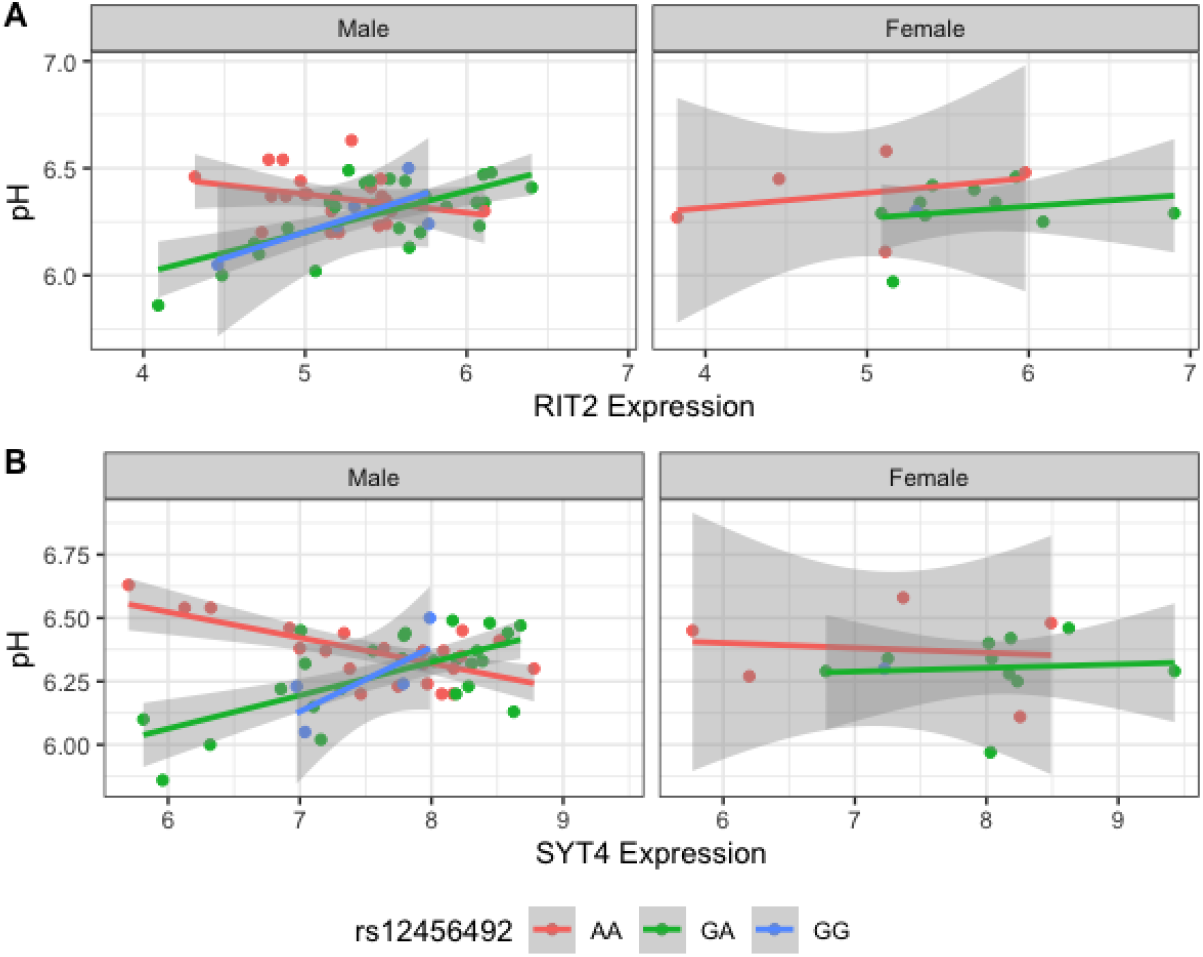
Scatter plot of pH and gene expression grouped by risk SNP genotype and stratified by sex within BrainEac sample for *RIT2* (A) and *SYT4* (B). Genotype groups are colored with lines representing linear fits with shaded areas marking the 0.95 confidence interval.

### Rs12456492 is not associated with *RIT2* and *SYT4* expression and pH within the substantia nigra in the GTEx dataset

Next, we used the GTEx dataset to test for replication of our findings from the BrainEac sample. We obtained gene expression data for *RIT2* and *SYT4* in the substantia nigra region, along with genotype data for the PD risk SNP rs12456492 yielding a sample size of 113. The pH ranged from 5.58 to 6.79, with a mean of 6.20. Similarly, we removed pH outliers from the samples using the same criteria as done with the BrainEAC dataset, resulting in 104 samples.

Correlation between gene expression of *RIT2* or *SYT4* with pH was significant (*RIT2*: r=0.27, p <0.006; *SYT4*: r=0.31, p<0.002). Next, we applied linear models on the GTEx data to first examine the influence of the risk SNP on pH level. The risk SNP by itself was not a significant predictor of pH (p = 0.14). Furthermore interaction between pH, rs12456492 and either *RIT2* or *STY4* was not observed (RIT2: t-stat = −1.10, p = 0.28, SYT4: t-stat = −0.84, p = 0.40). After accounting for the covariates (sex, age, RIN and PMI), the gene expression and SNP interactions were still not statistically significant terms in the models. Also, sex-specific signals were not observed in the GTEx substantia nigra samples.

### Shorter PMI values in GTEx

To explain the failed replication in the GTEx dataset, we examined the differences between the two cohorts. We noticed that PMI was longer in BrainEAC when compared to the GTEx samples. For the BrainEac, PMI ranged from 31 to 99 hours, whereas GTEx ranged from 4.78 to 23.13 hours. Hence there was no overlap between the two datasets as seen in Figure 5.

**Figure 5.**
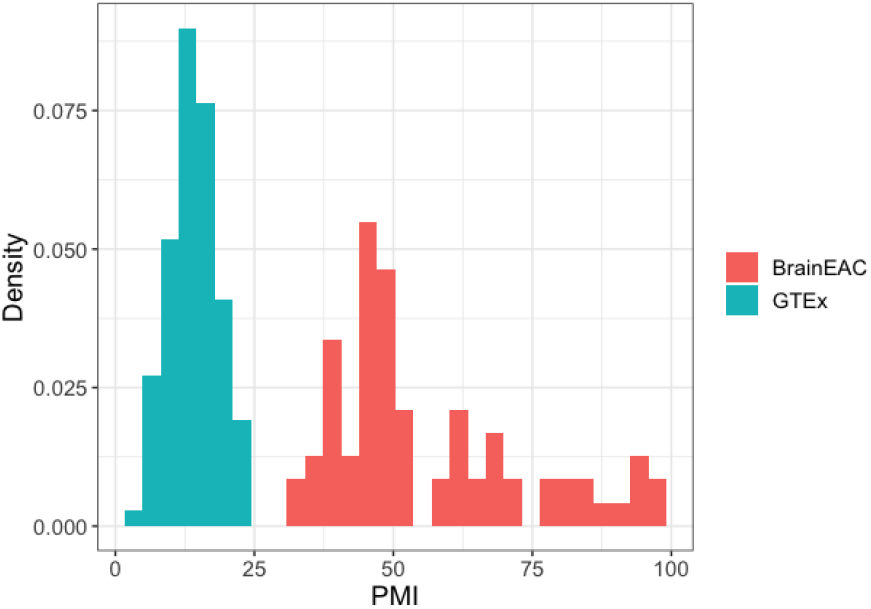
Density plot of PMI values in hours. GTEx and BrainEac samples are colored in blue and red, respectively.

### PMI influences pH-eQTL strength

To further investigate this difference, we stratified BrainEAC based on PMI. We split the sample based on the median PMI (49 hours). Using Model 2 and 3, interactions between gene expression and genotype were not statistically significant in short PMI (*RIT2*: p=0.15; *SYT4*:p=0.24) but were in long PMI (p<0.01 for both genes). The interaction between SNP, gene expression and PMI was not statistically significant for *RIT2* (B=-0.005 [-0.012 - 0.003], p=0.20), but was for *SYT4* (B= −0.004 [-0.008 - −0.0007], p<0.02). In Figure 6, G allele carriers in the stratified short and long PMI group show a positive correlation between gene expression and pH level. Although the long PMI group lacks GG carriers, a switch from positive to negative correlation is observed when comparing the GA and AA groups. This suggests the PMI difference between the datasets may explain why the pH-eQTL interaction was not observed in the GTEx sample.

**Figure 6.**
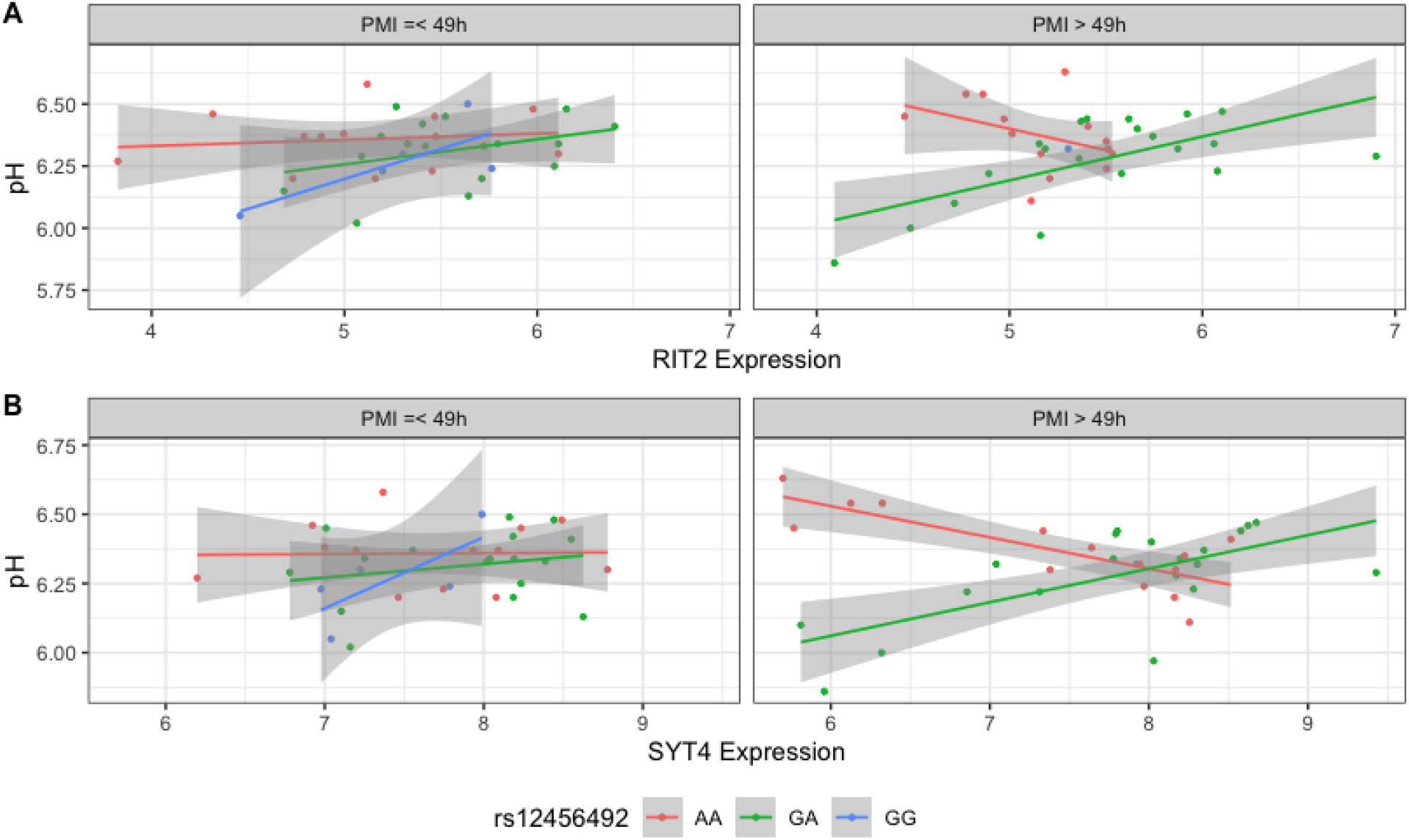
Scatter plot of pH and gene expression grouped by risk SNP genotype and stratified PMI within BrainEac sample for *RIT2* (A) and *SYT4* (B). Genotype groups are colored with lines representing linear fits with shaded areas marking the 0.95 confidence interval.

### *CA10* is co-expressed with *SYT4* and *RIT2*

Using a co-expression search tool, the two genes most co-expressed with *RIT2* were carbonic anhydrase 10 (*CA10*) and *SYT4*. Of the 97 brain datasets that excluded cancer studies, based on co-expression, the most relevant dataset was a study of Parkinson’s disease [GSE20146, (Zheng et al. 2010)]. In this top dataset that assayed expression in the globus pallidus interna (n=19), *RIT2* expression was correlated with *CA10* (r=0.52, p<0.03) and *SYT4* (r=0.75, p<0.0003), but *SYT4* was not co-expressed with *CA10* (r=0.43, p=0.065). We didn’t observe differential co-expression when comparing cases and controls. We note that CA10 gene expression is not a significant predictor for pH when accounting for the other covariates (sex, age, PMI, and RIN) in either BrainEAC or GTEx substantia nigra samples. Therefore the addition of *CA10* does not improve the models in predicting pH.

## Discussion

This study replicates the positive correlation between *RIT2* gene expression and pH in the human brain. Furthermore, we found that a co-localized PD associated genetic variant altered this relationship between expression and pH. When tested across ten brain regions, this interaction is specifically found in the substantia nigra, the primary location of Parkinson’s disease pathology. A similar association was found for the neighbouring *SYT4* gene. For carriers of the protective allele, the brain-wide positive correlation between gene expression and pH is inverted in the substantia nigra. In a combined model with both genes, the *SYT4* relationship is stronger. We attempted to replicate these findings using the GTEx dataset. However, the association between SNP interaction and gene expression of either *SYT4* and *RIT2* with pH was not seen. We observed that the PMI values were longer in the BrainEac cohort compared to the GTEx cohort. After stratifying the BrainEac data into short and long PMI, we only observed the relationship in the longer PMI group. These associations implicate interactions between rs12456492, *RIT2*, and *SYT4* in regulating pH in the brain.

When stratified by sex, the pH-eQTL relationship was not observed in females. This was more evident for *SYT4* than *RIT2*. The prevalence of PD is higher in males than females (Marras et al. 2018). Although our analysis was limited by sample size, we postulate that this sex-specific effect may help understand differences in PD incidence.

We suspect that we did not validate our findings in the GTEx dataset because of short PMI values in comparison to the BrainEAC samples. The differences between the datasets are clear, as there is no overlap between the PMI values. When the BrainEac cohort is split into short and long PMI groups, the results replicate the cohort differences. Specifically, the pH-eQTL relationship is observed in the samples with long but not short PMI. In agreement, studies of postmortem gene expression have found gene, and genotype-dependent associations with PMI (Zhu et al. 2017; Scott et al. 2020). Although we performed our analyses on neuropathologically normal brains, we speculate that a longer PMI value represents a neurodegenerative state that is closer to a Parkinsonian brain. Follow-up studies of postmortem samples or cell cultures derived from PD patients are warranted to test our findings in an experimental setting.

Several studies have examined pH-dependent interactions in the context of Parkinson’s disease. For example, α-synuclein aggregation and stability are increased at acidic pH (Lv et al. 2016; Pham et al. 2009). In addition, β-synuclein, an inhibitor of α-synuclein aggregation, is sensitive to pH and forms fibrils in mildly acidic pH (Moriarty et al. 2017; Williams et al. 2018). Using quantum chemical methods, Umek and colleagues determined that an acidic environment is required to prevent dopamine autoxidation (Umek et al. 2018). Kinetic modelling has also revealed that pH interactions with iron and dopamine could lead to oxidative stress (Sun et al. 2018). Caffeine consumption is associated with mild alkalosis (Tajima 2010) and a decreased risk of PD (Costa et al. 2010). We also note that *RIT2* is differentially co-expressed with interferon-gamma signalling genes in substantia nigra samples from PD cases (Liscovitch and French 2014). While indirect, interferon-gamma is acid-labile, and it’s overexpression in mice causes nigrostriatal neurodegeneration (Piasecki 1999; Chakrabarty et al. 2011). Mitochondria, which internally maintain an alkaline pH, are thought to be dysfunctional in PD (Chen, Turnbull, and Reeve 2019).

Mitochondrial dysfunction can lead to oxidative stress, resulting in lactic and intraneuronal acidosis (Arias et al. 2008; Koga et al. 2006; Balut et al. 2008). Recently, Rango and colleagues found that carriers of *PINK1* mutations, which are associated with early-onset PD, have abnormal pH levels in the visual cortex. Specifically, carriers of homozygous *PINK1* mutations had a higher pH at rest when compared to healthy controls and patients without *PINK1* mutations. Unlike healthy controls, pH did not increase upon activation in homozygous *PINK1* mutation carriers (Rango et al. 2020). Taken together, future studies of *RIT2* and *SYT4* that examine pH in the context of interferon-gamma, dopamine autoxidation, caffeine consumption, and mitochondrial function are warranted.

Both *SYT4* and Carbonic Anhydrase 10 (*CA10*) are co-expressed with *RIT2*. Unlike *RIT2* and *SYT4*, carbonic anhydrases are known to regulate intracellular and extracellular pH (Wingo et al. 2001). However, *CA10* has been found to be catalytically inactive and forms complexes with synaptic proteins (Sterky et al. 2017; Sjöblom et al. 1996; Nishimori et al. 2013). In agreement, *CA10* gene expression was not a predictor for brain pH when added to either the *RIT2* or *SYT4* models. Recently, Payan-Gomez and colleagues performed a co-expression network analysis of the human prefrontal cortex. In this analysis that used brain samples from old and young individuals, *CA10* was identified as a central gene in the network (Payán-Gómez et al. 2019). A related gene, *CA2*, was found to be elevated in mitochondria from middle-aged mouse brain samples (Pollard et al. 2016). Furthermore, carbonic anhydrase inhibitors have been found to prevent amyloid beta-induced mitochondrial toxicity (Solesio et al. 2018; Fossati et al. 2016). Further study of interactions between *RIT2* and *CA10* may reveal possible pH regulation mechanisms that are relevant to PD.

## Conclusion

This study described a relationship between gene expression and pH that interacted with genetic variation. Our analysis of this pH-eQTL relationship is constrained to the RIT2 locus that is strongly associated with PD risk and is only observed in substantia nigra. Additional interactions with sex and PMI were observed for SYT4 and, to a lesser degree, RIT2. These previously unknown associations suggest new mechanistic roles for rs12456492, RIT2, and SYT4 in the Parkinsonian brain.

## Acknowledgements

We thank the GTEx team for adding brain pH to the database. We thank Drs. Suneil Kalia and Lorraine Kalia for their insightful comments and suggestions.

This study was supported by the CAMH Foundation, CAMH Discovery Fund, and a National Science and Engineering Research Council of Canada (NSERC) Discovery Grant to LF.

